# Passenger co-deletion confounds glutaminolysis signatures anchored on *PTEN* loss: a cautionary case for location-aware signature design

**DOI:** 10.64898/2026.09.18.752756

**Authors:** Abdelmuhsen M. Abusneina, Soad M. Elwirfli

## Abstract

*PTEN*-deficient tumors are widely assumed to be glutamine-dependent, a view that has helped motivate the clinical development of glutaminase inhibitors. Whether glutaminolytic transcription scales with *PTEN* copy-number loss in human tumors, and by what mechanism, has not been established. We classified TCGA PanCancer Atlas tumors by GISTIC copy number as *PTEN* intact, hemizygous, or homozygous deletion, and scored a five-gene glutaminolysis signature (*GLS*, *SLC1A5*, *GOT1*, *GLUD1*, *GPT2*) against loss severity across fourteen tumor types, testing *MYC* mediation and chromosome-10 co-deletion as alternative mechanisms, with *GLS* dependence assessed in DepMap CRISPR data. The signature decreased with *PTEN* loss in all fourteen types, significantly in twelve, but was not *MYC*-mediated. The decline tracked chromosomal position rather than pathway membership: *GLUD1* and *GOT1* flank *PTEN* on 10q, and thirty-seven neighboring genes carrying no glutaminolysis annotation tracked *PTEN* copy number just as closely (mean rho 0.843 compared with 0.842), with co-deletion fidelity falling monotonically with distance from *PTEN* (rho = −0.993). Adjustment for genome-wide aneuploidy left the association intact, whereas conditioning on each gene’s own copy number abolished it for the chromosome-10 genes and left the others unchanged. In a natural experiment contrasting *PTEN* point-mutant, copy-neutral tumors with copy-neutral wild-type tumors, *GLUD1* was lower in the mutant group when all cohorts were pooled (P = 1.0 x 10^-8), an effect attributable to endometrial carcinoma, the one lineage in which mutation alone lowered *GLUD1*; excluding that lineage in a post hoc sensitivity analysis left no detectable difference (P = 0.22, equivalence P = 0.015), whereas hemizygous and homozygous deletion lowered *GLUD1* dose-dependently in every analysis. *GLS* dependence did not differ across dosage tiers. Prostate carcinoma was the sole departure in the negative direction, its off-chromosome-10 decline surviving adjustment for tumor purity, stage, *ERG* status and proliferation, opposite in direction to the signaling prediction; breast and stomach showed small positive associations. The inverse *PTEN*-glutaminolysis relationship in this transcript signature is therefore explained predominantly by passenger co-deletion rather than by a *PTEN*-linked metabolic program, making *PTEN* copy number unreliable as a stand-alone glutaminolysis biomarker and illustrating a general hazard: any signature scored against a frequently co-deleted locus can be confounded by where its genes sit. We provide a four-step screening procedure: decompose the signature by chromosomal position relative to the driver; test each constituent gene’s own copy number against the driver’s; compare driver-mutant copy-neutral tumors with copy-neutral wild-type tumors; and check the inferred pathway activity against an independent functional readout.

## Introduction

Proliferating tumor cells depend on glutamine to supply nitrogen for biosynthesis and to replenish tricarboxylic acid cycle intermediates through glutaminolysis, the committed step of which is catalyzed by glutaminase (*GLS*) [1]. Import depends on transporters such as *SLC1A5*, and downstream handling involves the aminotransferases *GOT1* and *GPT2* and the dehydrogenase *GLUD1*. Because glutaminase inhibitors have entered clinical testing [2], defining which tumors depend on this pathway is of therapeutic relevance.

Phosphatase and Tensin homolog (PTEN) is a lipid phosphatase that restrains Phosphoinositide 3-kinase (PI3K)/Protein Kinase B (AKT)/mammalian target of rapamycin complex 1 (mTORC1) signaling, and its loss is among the most common events in human cancer. In cell-line systems, mTORC1 activation after *PTEN* loss raises glutamine transporter expression and, through MYC, increases glutaminase. PTEN has separately been shown to promote the association of the anaphase-promoting complex with CDH1 [3], and APC/C-CDH1 to direct glutaminase for degradation [4], suggesting a further post-transcriptional route, although that link has not been demonstrated directly. From this work the field has generalized to the expectation that *PTEN*-deficient tumors are broadly glutamine-addicted. *PTEN* is also haploinsufficient, and subtle variation in its dose determines cancer susceptibility [5], so its effects are expected to be graded with gene dosage; it is inactivated in human tumors predominantly through copy-number loss that can be scored as one-copy or two-copy events.

Glutaminase has attracted sustained interest as a therapeutic target [6], and the expectation that *PTEN*-deficient tumors are glutamine-dependent has helped motivate the clinical development of glutaminase inhibitors. Yet the evidence linking *PTEN* specifically to glutaminolysis derives largely from individual cell-line models and from prostate cancer, and the two are not equivalent: a signaling relationship demonstrated in one lineage need not generalize, and an acute effect in cultured cells need not survive as a steady-state transcriptional difference in bulk tumors shaped by selection, stroma, and nutrient supply. No study has asked, across many tumor types and with attention to gene dosage, whether the glutaminolytic transcriptional program actually tracks *PTEN* copy-number loss, nor whether any such association reflects regulation or a structural artifact. Resolving this matters both mechanistically and for the use of *PTEN* status as a metabolic biomarker. These observations generate a specific, testable prediction: if *PTEN* loss drives glutaminolysis through graded signaling, a glutaminolysis transcriptional signature should rise monotonically from intact through hemizygous to homozygous *PTEN* loss. This prediction is not hypothetical: systemic elevation of *PTEN* in mice reduces glutamine uptake and suppresses glutaminolysis [7], so the converse expectation follows directly from the same axis. A complication is rarely considered. *PTEN* lies at chromosome 10q23.31, and *PTEN* deletions frequently remove a segment of neighboring sequence. Two of the canonical glutaminolysis genes, *GLUD1* at 10q23.2 and *GOT1* at 10q24.2, lie within this region, raising the possibility that any relationship between *PTEN* loss and the signature could reflect physical co-deletion rather than signaling. We therefore tested not only whether the signature tracks *PTEN* dosage across cancers, but also which mechanism, signaling or co-deletion, accounts for the association, using *MYC* as a candidate transcriptional mediator and the chromosomal location of the signature genes as a direct discriminator (Fig 1). More broadly, this case illustrates a structural hazard for any transcriptional signature scored against a frequently co-deleted locus, independent of the specific genes or pathway involved, and it motivates a general procedure for detecting such confounding.

**Fig 1.**
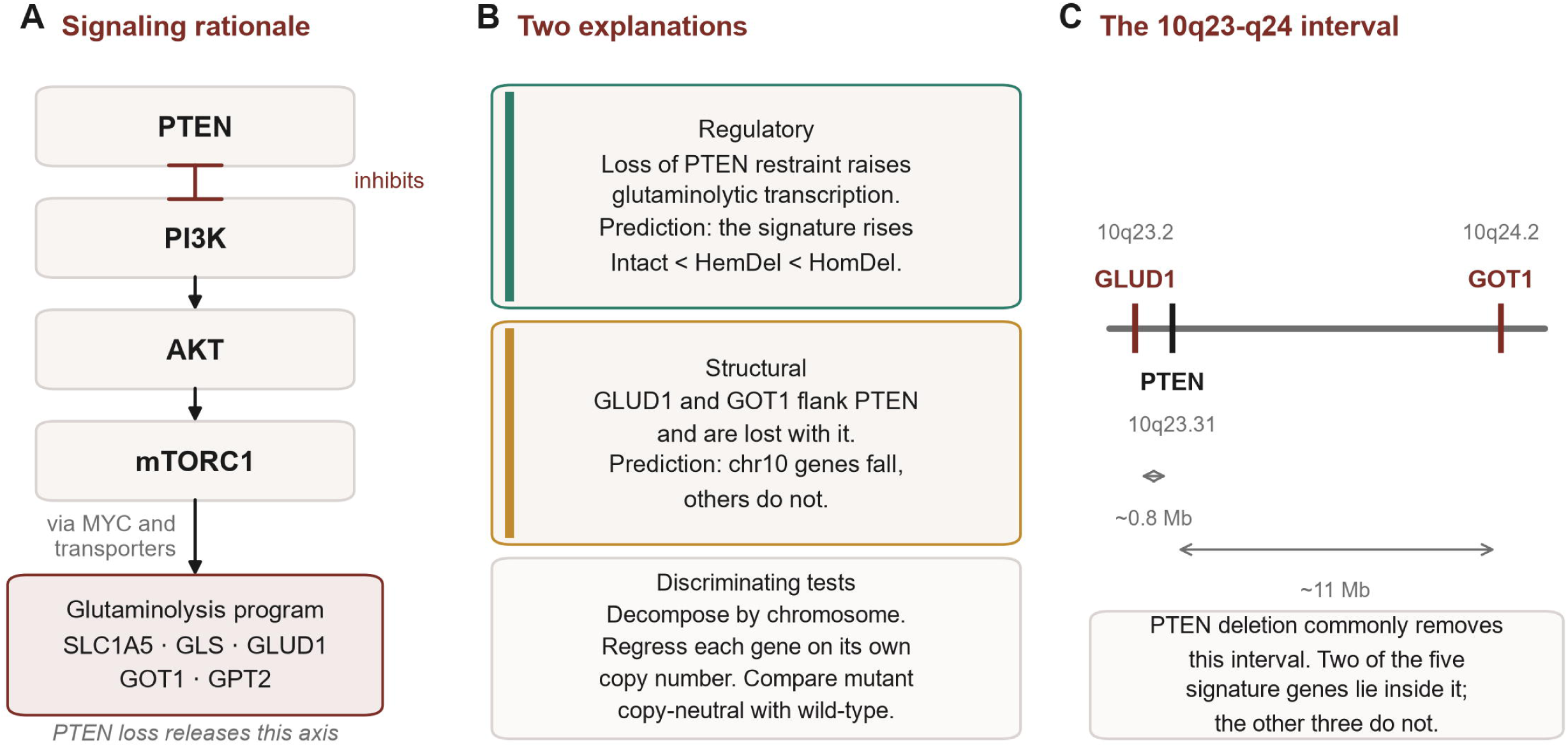
Competing explanations for an association between *PTEN* loss and the glutaminolysis signature. (A) PTEN restrains PI3K/AKT/mTORC1 signaling, which raises glutaminolytic gene expression through MYC and transporter induction; on this account *PTEN* loss should increase the signature. (B) The regulatory and structural explanations make opposite predictions, and three tests distinguish them. (C) Two of the five signature genes, *GLUD1* and *GOT1*, lie within the chromosome 10q23-q24 interval commonly removed by *PTEN* deletion; the remaining three lie on other chromosomes. Positions are indicative and drawn to relative scale within the interval rather than as a physical map.

## Materials and methods

### Data sources and cohorts

The Cancer Genome Atlas (TCGA) PanCancer Atlas RNA-seq expression (RSEM), Genomic Identification of Significant Targets in Cancer (GISTIC) copy number, mutation, and clinical data were obtained through the cBioPortal datahub [8,9]. Mutation data were retrieved through the cBioPortal REST API. Thirty-one solid-tumor PanCancer Atlas studies were assessed, one per cancer type; acute myeloid leukemia was omitted as a non-solid malignancy and diffuse large B-cell lymphoma was retained as a solid-tissue lymphoma. A study entered the analysis only if it contained at least ten tumors in each of the hemizygous and homozygous deletion tiers and at least ten intact tumors. This rule was fixed before the additional studies were analyzed, so that cohort membership is answerable as every study meeting the rule rather than as a selected set. Fourteen studies met it and seventeen did not; the excluded studies are listed with their tier counts in S5 Table. Cell-line CRISPR gene-effect (Chronos) scores and whole-genome-sequencing copy-number data were obtained from DepMap Public 26Q1 (CRISPRGeneEffect.csv and OmicsCNGeneWGS.csv) [10]. Consensus tumor purity estimates were taken from the TCGA PanCanAtlas ABSOLUTE tables.

Copy-number status was taken from the thresholded GISTIC2 [11] calls distributed in the cBioPortal file data_cna.txt, not from segmented log-ratio values. *PTEN* tiers were assigned directly from that call: homozygous, or deep, deletion for a value of −2 or below; hemizygous, or shallow, deletion for −1; and intact for 0 or above. Samples without a *PTEN* copy-number value were left unassigned. Expression values were log2-transformed after adding one and converted to z-scores within each cohort using the mean and standard deviation across all samples of that cohort, not against a diploid reference subset. Because both the exposure and the outcome are scaled within the same cohort, the choice of reference population does not affect the copy-number argument. Where a gene symbol appeared more than once in a matrix the first occurrence was retained, and sample identifiers were truncated to the fifteen-character TCGA sample barcode to align the expression, copy-number, mutation and clinical tables.

### Ethics declarations

This study used only publicly available, de-identified data from The Cancer Genome Atlas (TCGA) PanCancer Atlas and the DepMap cell-line repository. No new human participants were recruited, no new biospecimens were collected, and no new animal experiments were performed; accordingly, no additional institutional ethical approval or informed consent was required for this analysis.

### PTEN dosage, signature, and statistics

*PTEN* tiers were assigned from the GISTIC call as described above. The glutaminolysis signature comprised five genes selected a priori to represent the core enzymatic and transport steps of glutaminolysis rather than by data-driven selection: the glutamine transporter *SLC1A5*, glutaminase (*GLS*), and the enzymes handling glutamate and alpha-ketoglutarate, namely *GLUD1*, *GOT1*, and *GPT2*. The signature score was the mean of per-gene log2 z-scores across the five genes. Dose-response was tested by two-tailed Spearman correlation between an ordinal loss-severity score and the signature, with Benjamini-Hochberg correction across included tumor types; an adjusted P value below 0.05 was considered significant. Effect sizes are reported as median differences between tiers, as the variance explained by each correlation, and as Cliff’s delta with percentile bootstrap 95 percent confidence intervals from 2,000 resamples, computed for the signature and for each gene comparing *PTEN*-intact tumors with each deleted tier (Table 1, S2 Table). The ordering of the three tier medians is reported so that non-monotonic patterns are visible. *MYC* mediation was assessed by testing the association between loss severity and the mediator, using *MYC* expression and Hallmark MYC Targets V1 and V2 single-sample scores. Co-deletion was tested by contrasting chromosome-10 (*GLUD1*, *GOT1*) and off-chromosome-10 (*GLS*, *GPT2*, *SLC1A5*) sub-signatures, by correlating each gene’s copy number with *PTEN* copy number, and by extending the copy-number analysis to thirty-seven genes across 10q23-q24 carrying no glutaminolysis annotation, with positions taken from Ensembl (GRCh38). Distances are measured between transcription start sites, from each gene to that of *PTEN*, taking the start site of the Ensembl canonical transcript of each gene; genes with alternative first exons therefore have a single position, which may differ by a few hundred kilobases from the outermost transcript boundary. Every position used is deposited with the analysis code. To test whether the chromosome-10 association was independent of global aneuploidy, each correlation was recomputed as a Spearman partial correlation controlling for the per-sample aneuploidy score reported in the cBioPortal clinical data. Because a genome-wide score cannot isolate loss of a single chromosome arm, the analysis was repeated conditioning on a 10q arm-level proxy, defined as the median copy number of the interval genes lying more than 2 Mb from *PTEN*, and then on each gene’s own copy number. To separate loss of *PTEN* function from loss of the chromosomal segment, samples were classified by *PTEN* mutation status, counting non-silent variants as inactivating, with a truncating-only subset analyzed as a sensitivity check and a further analysis stratified by microsatellite instability and mutational burden. The primary pooled comparison includes all fourteen cohorts; a post hoc sensitivity analysis excluding uterine endometrial carcinoma is reported in the Results. Tumors carrying an inactivating *PTEN* mutation while retaining neutral or gained *PTEN* copy number were compared with copy-neutral *PTEN*-wild-type tumors and with hemizygously deleted tumors by two-tailed Mann-Whitney U test, per tumor type and pooled after within-cohort z-scoring; equivalence was assessed by two one-sided tests against a margin of 0.25 within-cohort standard deviations, fixed in advance. In prostate carcinoma the off-chromosome-10 composite was additionally modeled adjusting for tumor purity, pathological T stage, ERG expression and a proliferation index. DepMap *GLS* dependence was compared across tiers by two-tailed Kruskal-Wallis and Spearman tests. Summaries reported as means across cohorts weight every cohort equally, so that the largest cohorts do not dominate the pan-cancer picture. Analyses used Python with pandas, numpy, scipy, statsmodels, matplotlib and lifelines; code is provided in full. Reported cell-line gene-effect scores are Chronos estimates [12].

**Table 1.**
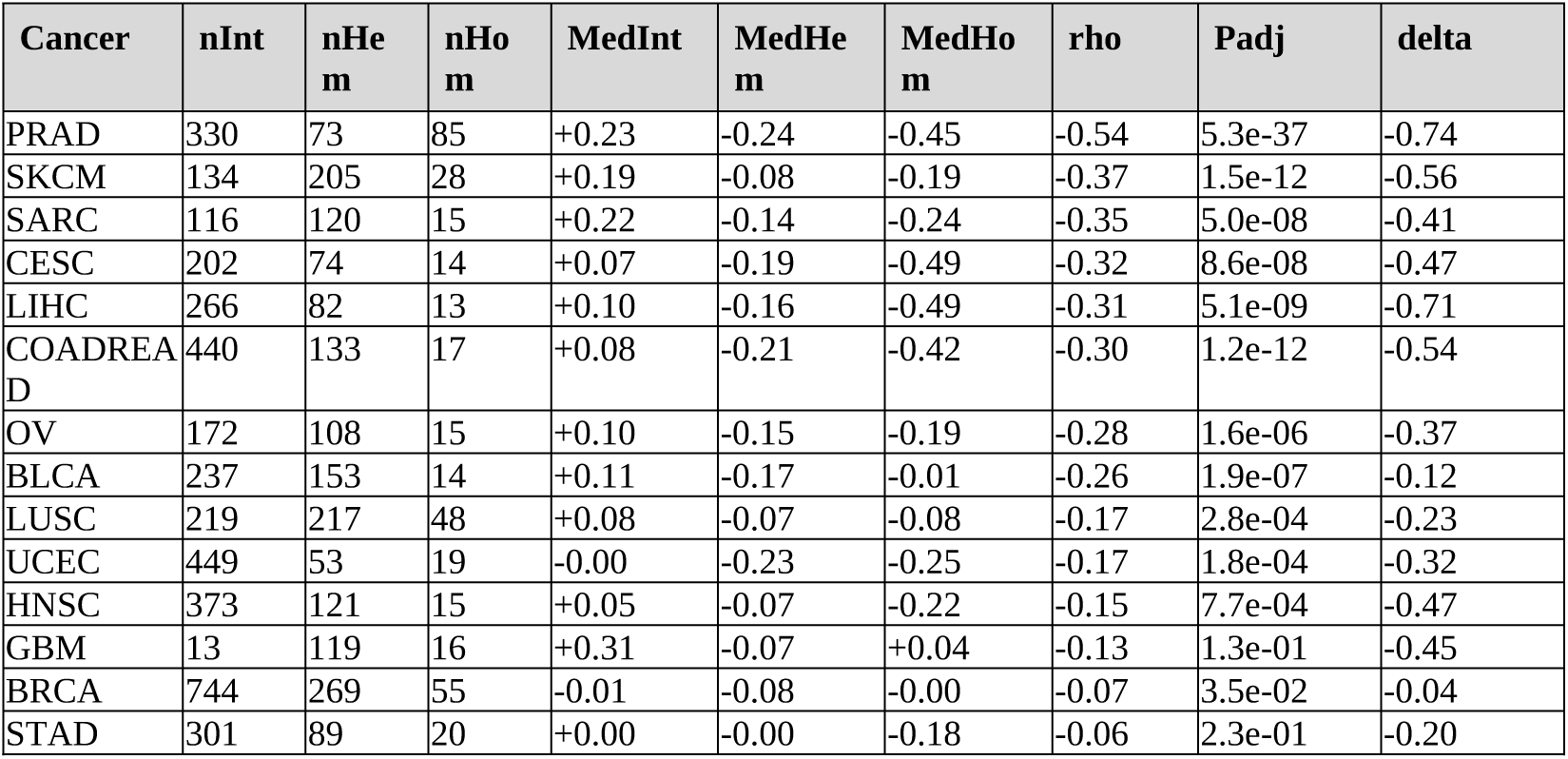
Glutaminolysis signature across *PTEN* copy-number dosage, ranked by trend strength. Medians are signature z-scores for the intact, hemizygous and homozygous tiers; rho is the Spearman correlation with loss severity; P is Benjamini-Hochberg adjusted across fourteen tumor types; delta is Cliff’s delta for the intact against homozygous comparison, with bootstrap confidence intervals given in S2 Table. nInt, nHem and nHom are the numbers of tumors in the intact, hemizygous and homozygous deletion tiers; MedInt, MedHem and MedHom are the corresponding median signature z-scores.

| Cancer | nInt | nHem | nHom | MedInt | MedHem | MedHom | rho | Padj | delta |
| --- | --- | --- | --- | --- | --- | --- | --- | --- | --- |
| PRAD | 330 | 73 | 85 | +0.23 | -0.24 | -0.45 | -0.54 | 5.3e-37 | -0.74 |
| SKCM | 134 | 205 | 28 | +0.19 | -0.08 | -0.19 | -0.37 | 1.5e-12 | -0.56 |
| SARC | 116 | 120 | 15 | +0.22 | -0.14 | -0.24 | -0.35 | 5.0e-08 | -0.41 |
| CESC | 202 | 74 | 14 | +0.07 | -0.19 | -0.49 | -0.32 | 8.6e-08 | -0.47 |
| LIHC | 266 | 82 | 13 | +0.10 | -0.16 | -0.49 | -0.31 | 5.1e-09 | -0.71 |
| COADREAD | 440 | 133 | 17 | +0.08 | -0.21 | -0.42 | -0.30 | 1.2e-12 | -0.54 |
| OV | 172 | 108 | 15 | +0.10 | -0.15 | -0.19 | -0.28 | 1.6e-06 | -0.37 |
| BLCA | 237 | 153 | 14 | +0.11 | -0.17 | -0.01 | -0.26 | 1.9e-07 | -0.12 |
| LUSC | 219 | 217 | 48 | +0.08 | -0.07 | -0.08 | -0.17 | 2.8e-04 | -0.23 |
| UCEC | 449 | 53 | 19 | -0.00 | -0.23 | -0.25 | -0.17 | 1.8e-04 | -0.32 |
| HNSC | 373 | 121 | 15 | +0.05 | -0.07 | -0.22 | -0.15 | 7.7e-04 | -0.47 |
| GBM | 13 | 119 | 16 | +0.31 | -0.07 | +0.04 | -0.13 | 1.3e-01 | -0.45 |
| BRCA | 744 | 269 | 55 | -0.01 | -0.08 | -0.00 | -0.07 | 3.5e-02 | -0.04 |
| STAD | 301 | 89 | 20 | +0.00 | -0.00 | -0.18 | -0.06 | 2.3e-01 | -0.20 |

### Use of generative AI

All study design, analytical decisions, and interpretation of the findings are the authors’ own. The generative artificial intelligence assistant Claude (Anthropic) was used under author supervision to write and run the Python analysis code, and to assist with writing and language editing of the manuscript. The authors assume full responsibility for the accuracy and integrity of the work.

## Results

### *The glutaminolysis signature decreases with PTEN copy-number* loss across cancer types

Fourteen tumor types met the pre-specified inclusion rule of at least ten tumors in each dosage tier: prostate (PRAD), melanoma (SKCM), sarcoma (SARC), cervical (CESC), hepatocellular (LIHC), colorectal (COADREAD), ovarian (OV), bladder (BLCA), squamous lung (LUSC), uterine endometrial (UCEC), head and neck (HNSC), glioblastoma (GBM), breast (BRCA), and stomach (STAD). The remaining seventeen studies were excluded, in every case because of too few tumors in one or more dosage tiers, most often homozygous deletion; the full list with tier counts is given in S5 Table.

In every included type the glutaminolysis signature decreased as *PTEN* loss increased (Table 1, Fig 2), with a significant monotonic trend in twelve after correction and the strongest effect in prostate cancer (Spearman rho = −0.54, adjusted P = 5.3 x 10^-37). No tumor type showed the positive trend predicted by the signaling model (Fig 3). Effect sizes varied widely: the association explained 29.1 percent of variance in prostate but 0.5 percent in breast and 0.4 percent in stomach, where correlations of −0.07 and −0.06 are detectable only because of cohort size. Cliff’s delta for the intact against homozygous comparison tells the same story, from −0.74 in prostate and −0.71 in hepatocellular carcinoma to −0.04 in breast, whose confidence interval includes zero (Table 1; per-gene values with intervals in S2 Table). In three cohorts, bladder, glioblastoma and breast, the three tier medians were not monotonically ordered, so the negative coefficient in those cohorts summarizes a trend that is not consistently graded. Because glioblastoma had only thirteen *PTEN*-intact tumors, an unstable baseline for a tumor type in which *PTEN* loss is nearly ubiquitous, its estimate should be interpreted with particular caution.

**Fig 2.**
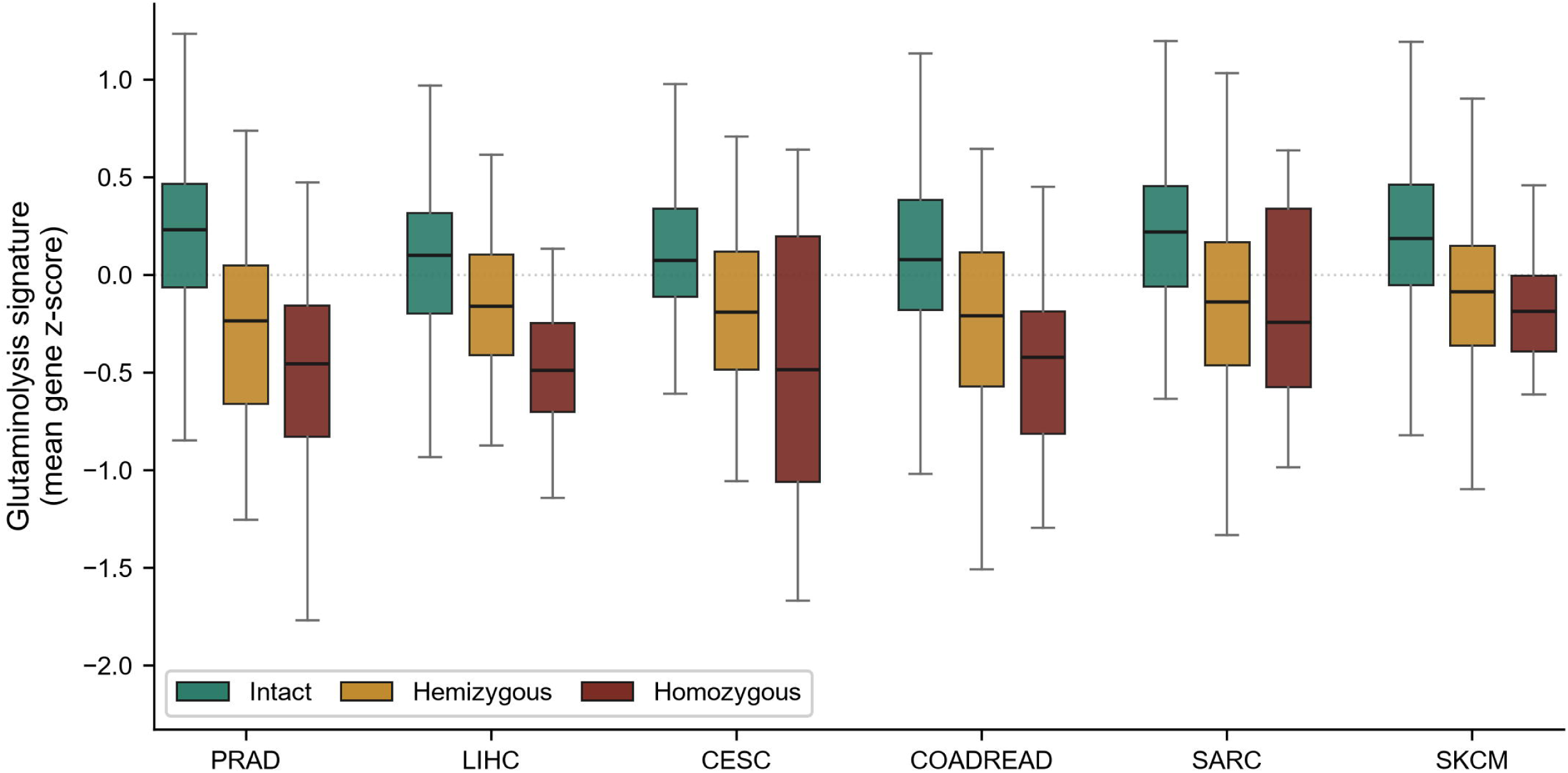
Glutaminolysis signature by *PTEN* dosage in the six tumor types with the largest decline from intact to homozygous deletion. The signature declines from intact through hemizygous to homozygous deletion. The remaining eight cohorts are shown in S2 Fig.

**Fig 3.**
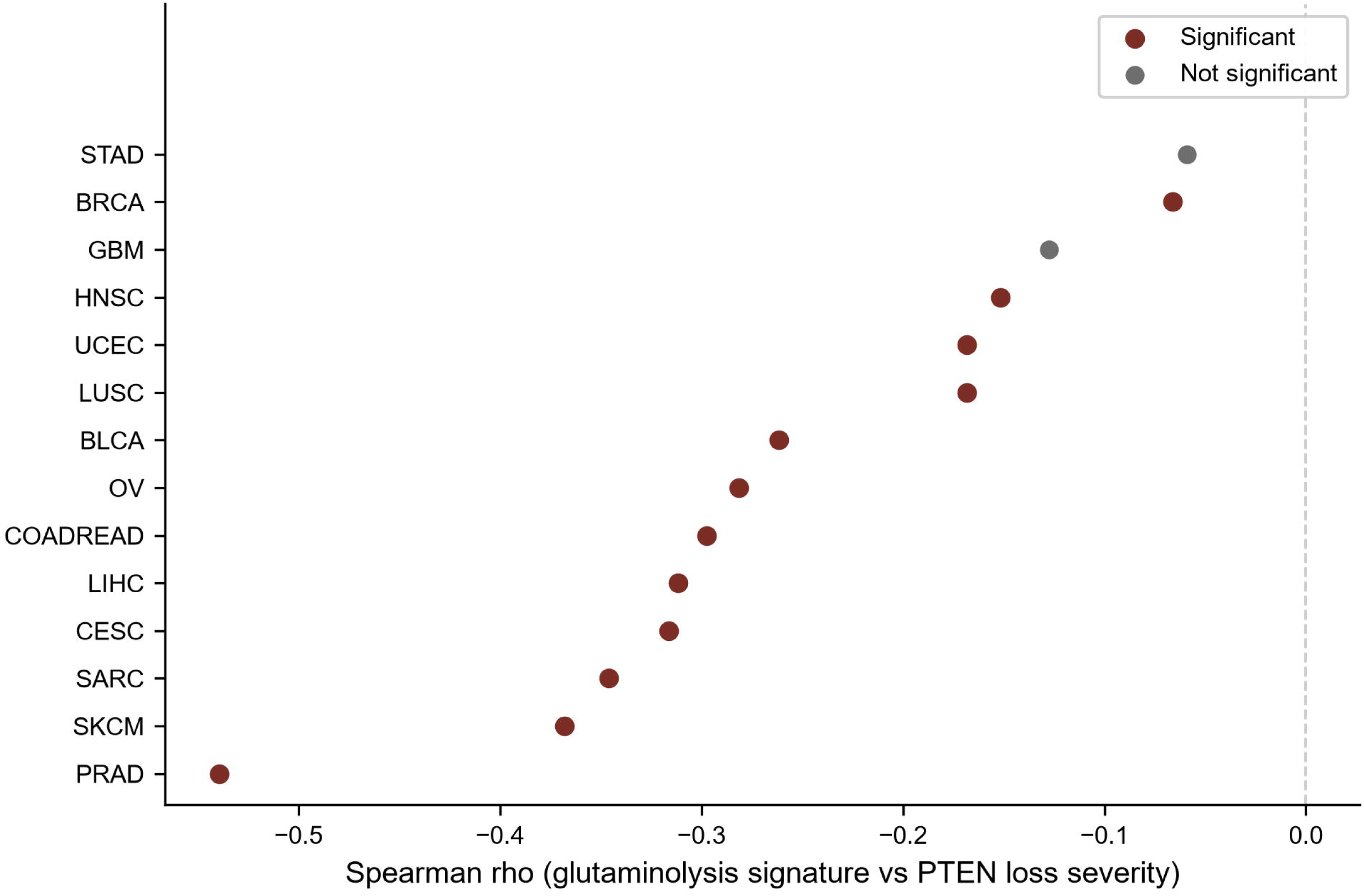
Dose-response across all fourteen included tumor types, ordered by effect size. All trends are negative; markers in dark red indicate types significant after Benjamini-Hochberg correction, gray markers those that are not.

### *Co-deletion is a property of the chromosomal interval, not of the* pathway

If *GLUD1* and *GOT1* fall with *PTEN* loss because of a metabolic relationship, then genes in the same chromosomal neighborhood with no connection to glutaminolysis should not behave the same way. We therefore extended the analysis to thirty-seven genes across 10q23-q24 carrying no glutaminolysis annotation (S6 Table), taking positions from Ensembl (GRCh38) and correlating each gene’s copy number with *PTEN* copy number in every cohort.

The unannotated neighbors tracked *PTEN* copy number as closely as the signature genes did. Across fourteen cohorts the mean correlation was 0.843 for the thirty-seven unannotated genes and 0.842 for *GLUD1* and *GOT1*, while the three signature genes on other chromosomes averaged 0.034. Co-deletion fidelity declined monotonically with distance from *PTEN* (Spearman rho between distance and correlation = −0.993; Fig 4). Because neighboring genes share copy-number profiles, a nominal P value computed as though the genes were independent observations would overstate the evidence, and none is reported. *GLUD1*, at 0.77 Mb from the *PTEN* transcription start site, reached 0.891, which is lower than *KLLN* (0.965), *ATAD1* (0.954) and *PAPSS2* (0.937), each of which lies within 0.3 Mb of *PTEN*, and indistinguishable from *RNLS* (0.948), which lies at a comparable distance. Fidelity falls further in *FAS* (0.908) and *ACTA2* (0.910), which lie beyond 1 Mb. *GOT1*, at 11.57 Mb, reached 0.793, between HPSE2 at 11.37 Mb (0.800) and ABCC2 at 11.92 Mb (0.787). Across the interval, co-deletion fidelity for the two signature genes fell within the range spanned by the unannotated neighbors at comparable distances.

**Fig 4.**
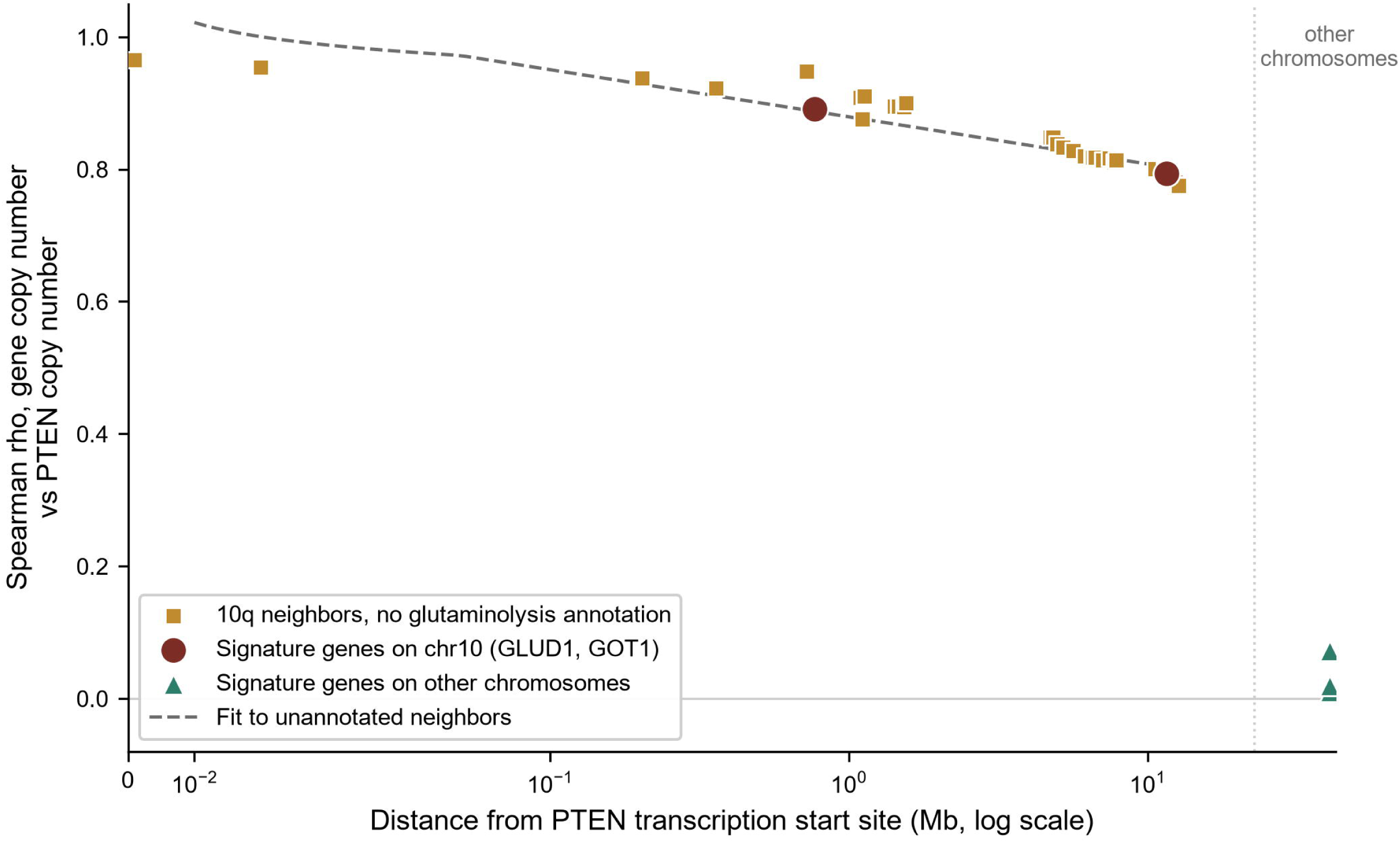
Co-deletion with *PTEN* declines with chromosomal distance and is unrelated to pathway membership. Each point is one gene, showing the mean Spearman correlation between its copy number and *PTEN* copy number across fourteen tumor types, plotted against distance from the *PTEN* transcription start site. Gene positions are from Ensembl (GRCh38) and are deposited with the analysis code.

The deletions involved are focal rather than arm-level. Across fourteen cohorts the median span of co-deleted sequence in *PTEN* homozygously deleted tumors was 0.72 Mb, and 90 percent of such tumors carried deletions confined within 2 Mb of *PTEN*. That span is marginally shorter than the 0.77 Mb separating *PTEN* from *GLUD1*, so *GLUD1* sits at the outer edge of the typical focal homozygous event and tracks *PTEN* mainly through the broader, predominantly hemizygous 10q loss that accompanies it. *GOT1*, at 11.57 Mb, lies beyond all but the largest of those events, which is why its fidelity is lower again.

### Aneuploidy adjustment and gene-level copy-number adjustment give different answers

Two adjustments distinguish passenger co-deletion from a signaling effect. First, the correlation between *PTEN* loss severity and each signature gene was recomputed as a Spearman partial correlation controlling for the per-sample aneuploidy score (S3 Table). The *GLUD1* and *GOT1* associations were essentially unchanged, for example *GLUD1* partial rho −0.42 in breast, −0.55 in ovarian and −0.59 in prostate against unadjusted values of −0.45, −0.53 and −0.59, confirming that the signal is not an artifact of global ploidy.

A genome-wide aneuploidy score cannot, however, isolate loss of a single chromosome arm: a tumor with a focal 10q deletion and an otherwise quiet genome carries a low score. We therefore repeated the analysis conditioning first on a 10q arm-level proxy and then on each gene’s own copy number, which is the direct test. Conditioning on arm-level status removed most but not all of the association, moving the mean chromosome-10 coefficient from −0.321 to −0.081, as expected if the events that carry *GLUD1* are smaller than the arm. Conditioning on each gene’s own copy number abolished it, reducing the mean coefficient to −0.022, while the off-chromosome-10 genes were unaffected throughout, their mean coefficient moving from +0.012 unadjusted to −0.001 and +0.021 under the two adjustments. The two adjustments therefore differ in what they resolve: the genome-wide aneuploidy score does not separate the chromosome-10 genes from the others, whereas gene-level copy number does.

Separating the signature into its chromosome-10 and off-chromosome-10 components gave the same answer (Table 2, Fig 5). The chromosome-10 genes decreased with *PTEN* loss in all fourteen types (rho −0.12 to −0.56), whereas the off-chromosome-10 genes were flat or slightly positive, with a mean difference of −0.42. At the level of copy number the separation is absolute: *GLUD1* copy number correlated with *PTEN* copy number between 0.71 and 0.98 across tumor types (mean 0.89) and *GOT1* between 0.55 and 0.91 (mean 0.79), whereas the off-chromosome-10 genes showed no such relationship (*GLS* mean 0.01, *GPT2* 0.07, *SLC1A5* 0.02). Full per-gene results are given in S1 and S2 Tables.

**Fig 5.**
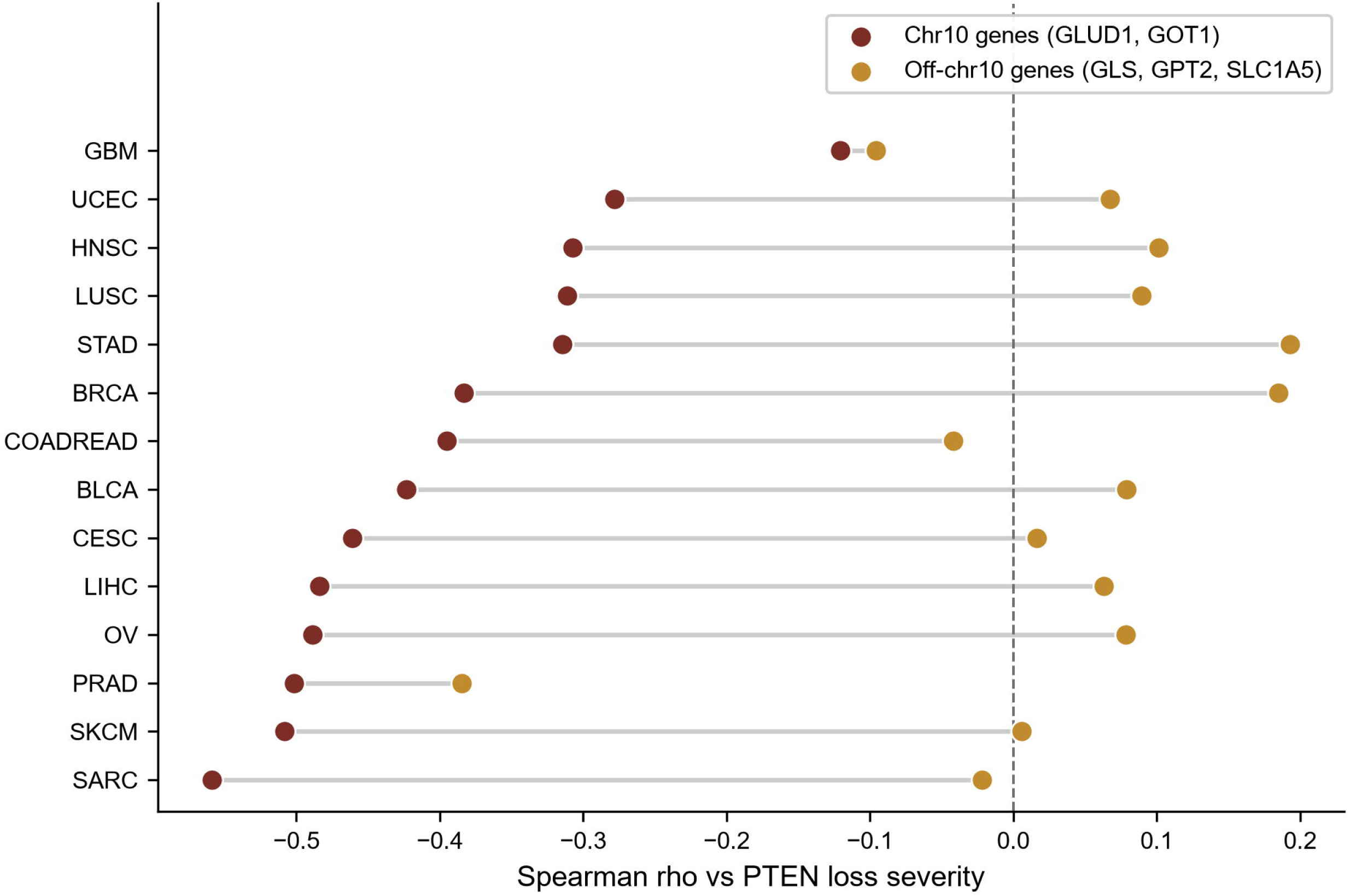
Decomposition of the signature by chromosomal location. Chromosome-10 signature genes (*GLUD1*, *GOT1*) fall with *PTEN* loss, while off-chromosome-10 genes (*GLS*, *GPT2*, *SLC1A5*) do not. Each line pairs the two sub-signatures within a tumor type.

**Table 2.** Contribution of chromosome-10 versus off-chromosome-10 signature genes to the association with *PTEN* loss severity. Values are Spearman rho against *PTEN* loss severity.

| Cancer | Chr10 genes rho | Off-chr10 genes rho | Difference |
| --- | --- | --- | --- |
| SARC | -0.56 | -0.02 | -0.54 |
| SKCM | -0.51 | +0.01 | -0.51 |
| PRAD | -0.50 | -0.38 | -0.12 |
| OV | -0.49 | +0.08 | -0.57 |
| LIHC | -0.48 | +0.06 | -0.55 |
| CESC | -0.46 | +0.02 | -0.48 |
| BLCA | -0.42 | +0.08 | -0.50 |
| COADREAD | -0.40 | -0.04 | -0.35 |
| BRCA | -0.38 | +0.18 | -0.57 |
| STAD | -0.31 | +0.19 | -0.51 |
| LUSC | -0.31 | +0.09 | -0.40 |
| HNSC | -0.31 | +0.10 | -0.41 |
| UCEC | -0.28 | +0.07 | -0.35 |
| GBM | -0.12 | -0.10 | -0.02 |

### A mutation-based natural experiment

To separate loss of *PTEN* function from loss of the chromosomal segment, tumors in which *PTEN* was inactivated by point mutation or small indel while retaining neutral or gained copy number were compared with copy-neutral *PTEN*-wild-type tumors, and with hemizygously deleted tumors as a positive control (S4 Table). Pooled across all fourteen cohorts, and reported here as the primary analysis, *GLUD1* expression in the mutant copy-neutral group was lower than in wild-type tumors (435 against 3,561 tumors, difference −0.24, P = 1.0 x 10^-8). Stratification showed this to be driven by endometrial carcinoma, the one lineage in which mutation alone lowered *GLUD1*: excluding it in a post hoc sensitivity analysis, the mutant copy-neutral group did not differ detectably from wild-type (mean z-score 0.23 against 0.31, difference −0.08, 90 percent CI −0.21 to +0.05, P = 0.22), and equivalence within a pre-specified margin of 0.25 standard deviations was supported (two one-sided tests, P = 0.015). Hemizygous deletion lowered *GLUD1* markedly (mean −0.44) and homozygous deletion further still (mean −0.89). Within individual cohorts other than endometrial carcinoma no difference was detected, including colorectal (P = 0.91), lung squamous (P = 0.07), breast (P = 0.85) and stomach (P = 0.54); these per-cohort groups are small, between one and twenty-five tumors outside endometrial carcinoma, so they establish the absence of a detectable effect rather than equivalence. A sensitivity analysis restricted to truncating variants gave the same result (difference −0.09, P = 0.36).

Endometrial carcinoma was the exception. In this hypermutated, *PTEN*-mutation-rich lineage (mutant, copy-neutral n = 320) *GLUD1* did fall with mutation alone (difference −0.43, P = 1 x 10^-5), and this single cohort accounts for the pooled signal when included, so it is reported separately. Because a hypermutated background could in principle generate passenger mutations without a mechanism, we repeated the comparison in microsatellite-stable tumors only. The effect was larger rather than smaller (n = 207 against 116, difference −0.53, P = 1.2 x 10^-6), and the same held when restricting to tumors below a mutational burden threshold (difference −0.55, P = 8.7 x 10^-6). Microsatellite instability therefore does not account for the endometrial exception.

### MYC does not mediate the association

Because MYC is the principal transcriptional driver of glutaminolytic genes [13], we tested whether it mediates the inverse association. Mediation requires an association between exposure and mediator. *PTEN* loss severity was not consistently related to *MYC* expression: the coefficient was significant in three of fourteen tumor types and the direction was inconsistent, negative in ten cohorts and positive in four. Because *MYC* messenger RNA is an imperfect proxy for MYC transcriptional activity, we repeated the analysis using Hallmark MYC Targets V1 and V2 single-sample scores, with the same result (significant in three and five of fourteen respectively, again in inconsistent directions; Fig 6). A mediator whose relationship to the exposure differs in sign between lineages cannot account for a consistent association. The *MYC* transcript and target-score measures tested here therefore provide no evidence of consistent mediation, although they do not exclude post-transcriptional *MYC* regulation that these measures cannot capture.

**Fig 6.**
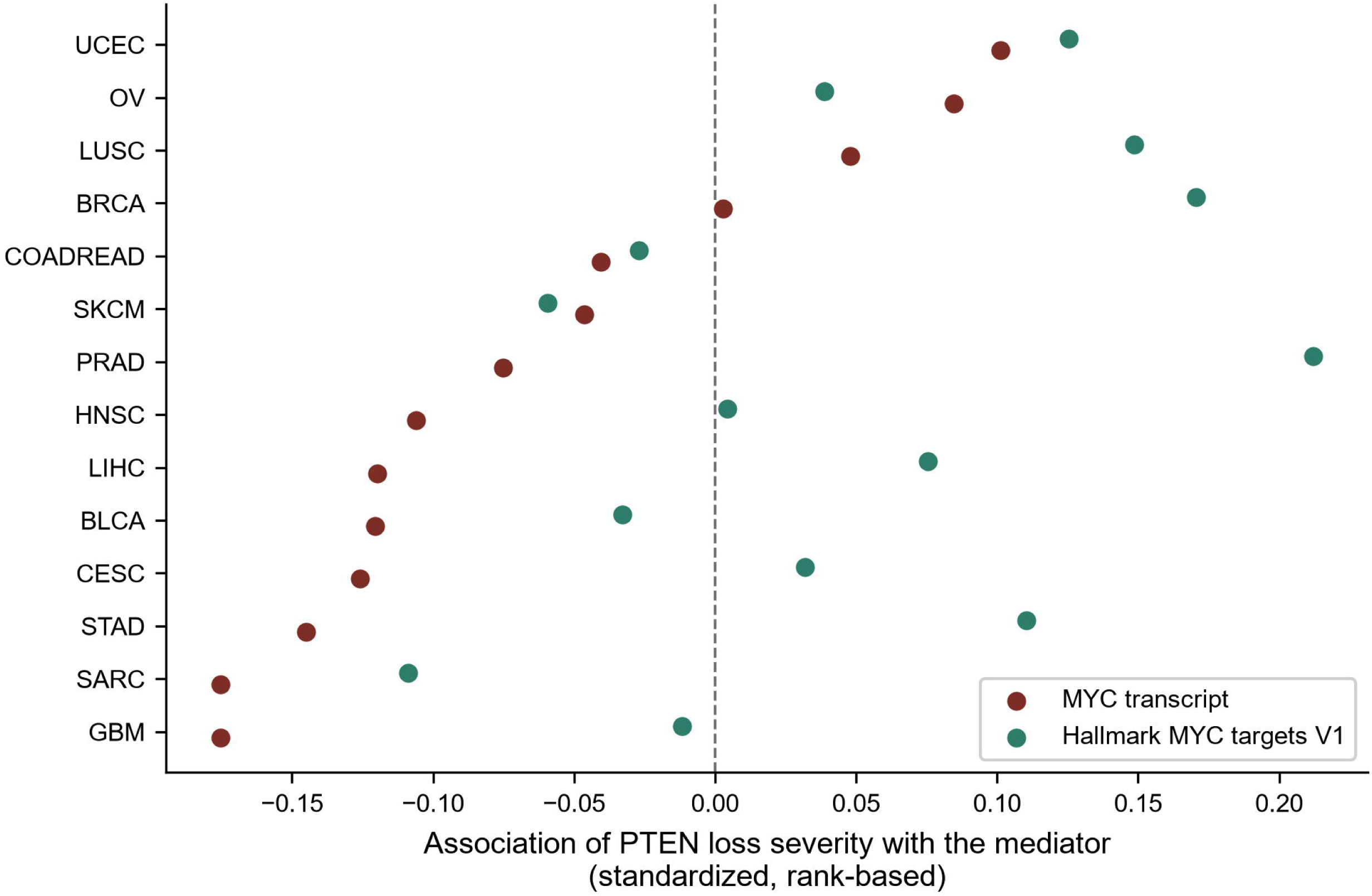
MYC does not mediate the inverse *PTEN*-glutaminolysis association. Association of *PTEN* loss severity with *MYC* transcript and with Hallmark MYC target scores, by tumor type. The direction is inconsistent between lineages.

### GLS dependence does not track PTEN dosage

If *PTEN* loss created a functional glutaminolytic dependency, *PTEN*-deficient cells should rely more on glutaminase. In DepMap CRISPR data, copy number was available for 858 of the 1,208 cell lines carrying a gene-effect score. *GLS* gene-effect scores did not differ across *PTEN* dosage tiers (median −0.147 intact, −0.135 hemizygous, −0.097 homozygous; Kruskal-Wallis P = 0.74; Spearman rho = 0.0008, P = 0.98), and equivalence within a margin of 0.15 Chronos units was supported (P = 0.003).

This result bounds rather than proves the absence of a dependency. *GLS* falls below the conventional dependency threshold of −0.5 in only 10.3 percent of cell lines, with a median gene-effect score of −0.15, whereas the common essential genes RPL23A and PSMA1 fall below that threshold in 100 percent of lines with median scores of −2.43 and −2.36. *GLS* is therefore a weak dependency in most lines irrespective of *PTEN* status, so the absence of a difference across tiers constrains the signaling model only weakly and is best read as corroborating the transcriptional and copy-number findings. Two-dimensional cultures in nutrient-rich media may also not reproduce the metabolic selection pressures of tumors in vivo.

### Prostate carcinoma departs from the passenger account

In thirteen of the fourteen cohorts the off-chromosome-10 genes were flat or slightly positive. After Benjamini-Hochberg correction across fourteen cohorts, three associations remained significant: breast (rho = +0.18, adjusted P = 8.3 x 10^-9) and stomach (rho = +0.19, adjusted P = 4.0 x 10^-4), both in the direction the signaling model predicts, and prostate (rho = −0.38, adjusted P = 1.7 x 10^-17), in the opposite direction. The breast and stomach effects are small, explaining three and four percent of variance respectively.

The prostate association is not explained by co-deletion, since *GPT2* and *SLC1A5* are not co-deleted with *PTEN* (copy-number correlations 0.05 and −0.07) and their expression associations are unchanged by conditioning on their own copy number (−0.33 to −0.32 and −0.35 to −0.36). Prostate also has the lowest co-deletion fidelity of the fourteen cohorts (*GLUD1* 0.77, *GOT1* 0.55, against means of 0.89 and 0.79), consistent with the characteristically focal nature of *PTEN* deletion in this tissue, so extensive passenger loss cannot account for it either.

We therefore tested whether the prostate association reflects features that co-vary with *PTEN* deletion in this tissue, adjusting for tumor purity, pathological T stage, ERG expression as a surrogate for TMPRSS2-ERG fusion, and a proliferation index. The association attenuated by 16.1 percent and remained clearly significant (standardized coefficient −0.385 to −0.323, P = 1.3 x 10^-11, n = 464), while the chromosome-10 composite attenuated by 10.7 percent as expected for a structural effect. At gene level the adjustment removed 52.9 percent of the *SLC1A5* association and 29.2 percent of *GPT2*, against 9.9 percent for *GLUD1* and 11.8 percent for *GOT1*, so roughly half of the *SLC1A5* effect is attributable to composition, stage, ERG or proliferation and the remainder is not.

In prostate carcinoma, then, the reduction in glutaminolytic transcription persists after adjustment for passenger co-deletion, tumor composition, stage, *ERG* status and proliferative activity. Its basis is not established by these data.

### *PTEN loss is prognostic independently of the glutaminolytic* program

Because *PTEN* loss is an established adverse prognostic marker, we examined whether *PTEN* dosage predicted overall survival in the included cohorts, and whether the glutaminolysis signature carried independent prognostic weight (S1 Fig). Increasing *PTEN* loss severity was associated with worse overall survival in uterine endometrial carcinoma and glioblastoma, and the glutaminolysis signature was not consistently prognostic. The prognostic effect of *PTEN* loss is therefore real in specific tissues but is not explained by, and does not track, the glutaminolytic signature. Independently of the signature, hemizygous *PTEN* loss has been linked to tumor immune evasion and worse outcome in pan-cancer analyses [14], consistent with a broader role for *PTEN* in maintaining immune surveillance that is mechanistically distinct from the co-deletion artifact described here [15].

## Discussion

The expectation that *PTEN* loss up-regulates glutaminolysis, extrapolated from PI3K/AKT/mTOR signaling, is not borne out in human tumor transcriptomes. Across fourteen tumor types the glutaminolysis signature decreased with *PTEN* loss, and direct testing showed this to be a consequence of gene location rather than of metabolic regulation. Two of the five signature genes, *GLUD1* and *GOT1*, lie within the chromosome 10q23-q24 interval that *PTEN* deletion commonly removes, and they behave exactly as any other gene in that interval behaves: their copy number tracks *PTEN* copy number with a fidelity determined by distance and nothing else.

The confound is not detected by the control most naturally applied to it. Adjusting for genome-wide aneuploidy left the chromosome-10 associations essentially unchanged, which would ordinarily be read as evidence against a structural explanation. A genome-wide score cannot isolate loss of a single arm: a tumor carrying a focal 10q deletion in an otherwise quiet genome scores low on aneuploidy while having lost precisely the segment in question. Conditioning instead on each gene’s own copy number abolished the association for *GLUD1* and *GOT1* and left the off-chromosome-10 genes untouched. The distinction matters beyond this locus, because aneuploidy adjustment is widely applied where gene-level adjustment is the appropriate test.

The positional basis of the association can be demonstrated directly. Extending the analysis to thirty-seven genes across 10q23-q24 with no annotated role in glutaminolysis, co-deletion with *PTEN* was as strong for those genes as for the two signature genes, and declined monotonically with distance from *PTEN*. *GLUD1*, at 0.77 Mb, was co-deleted less faithfully than three unannotated genes lying closer to *PTEN*, and indistinguishably from a fourth at a comparable distance. The deletions involved are focal: across fourteen tumor types the median span of co-deleted sequence was 0.72 Mb, and 90 percent of homozygously deleted tumors carried deletions confined within 2 Mb of *PTEN*. Whether a gene tracks *PTEN* copy number is therefore governed by where it sits on chromosome 10 rather than by what pathway it belongs to.

This interpretation reconciles a fragmented literature. *PTEN* controls metabolism substantially through post-transcriptional routes [16] that a transcriptional signature cannot capture: PTEN dephosphorylates AKT to limit GLUT1 surface expression and glucose uptake [17], promotes the association of the anaphase-promoting complex with CDH1 [3], which in turn can direct glutaminase for degradation [4], although that particular link has not been demonstrated directly, and is itself regulated by phosphorylation, which can impose a tumor-promoting metabolic state [18], and by deubiquitinating enzymes that tune its metabolic output [19,20]. *PTEN* loss also confers resistance to ferroptosis by increasing xCT-mediated cystine import through AKT-GSK3β-NRF2 signaling [21], and other tumor suppressors regulate glutaminolysis independently of *PTEN* copy-number status, as when NDRG2 restrains glycolysis and glutaminolysis by repressing c-Myc [22]; both observations reinforce that *PTEN*’s metabolic footprint is heterogeneous and only partly transcriptional. Reports that *PTEN* loss increases glutamine utilization describe flux-level phenomena, including routing of glutamine into de novo pyrimidine synthesis [23], rather than transcriptional up-regulation of glutaminolysis. Other lineages show that *PTEN* loss can reduce or redirect glutaminolysis: in T-cell acute leukemia *PTEN* loss rewires metabolism toward ATP-citrate-lyase-dependent lipogenesis [24], while Notch1-driven T-ALL cells separately depend on glutaminolysis for resistance to Notch pathway inhibition [25]; indeed, *PTEN* deletion alone is insufficient to drive the leucine-transport-dependent metabolic program of T-ALL, which instead requires NOTCH signaling [26], further indicating that glutamine dependence in this lineage is shaped by multiple, only partly *PTEN*-related, drivers. Against this background, the present finding adds that in most solid tumors the transcriptional appearance of reduced glutaminolysis is not regulatory at all but structural, arising from passenger co-deletion.

One lineage does not fit the passenger account. In prostate carcinoma the off-chromosome-10 genes also declined with *PTEN* loss, and this was the only such association among fourteen cohorts to survive correction. It is not explained by co-deletion, since *GPT2* and *SLC1A5* are not co-deleted with *PTEN* and their associations are unchanged by conditioning on their own copy number; nor by extensive passenger loss, since prostate shows the lowest co-deletion fidelity of the fourteen cohorts, consistent with the characteristically focal *PTEN* deletions of this tissue; nor by tumor composition, stage, ERG status or proliferative activity, which together account for 16 percent of the association and leave the remainder intact. The direction is nonetheless opposite to that predicted by release of PI3K/AKT/mTORC1 restraint, so it is not evidence of the signaling relationship described in this tissue [27,28,29]. This prostate association was examined after the pan-cancer result was in hand and is reported as exploratory. The basis of this residual association is unresolved. What can be said is that it survives exclusion of copy number, tumor composition, stage, ERG status and proliferative activity, so it is independent of the passenger co-deletion effect that accounts for the association in the other thirteen cohorts. Endometrial carcinoma showed a distinct pattern: *PTEN* inactivation by mutation alone lowered *GLUD1*, and this persisted in microsatellite-stable tumors, excluding hypermutation as its cause; again the direction is a reduction where signaling predicts an increase. Two further cohorts, breast and stomach, showed small associations in the direction the signaling model predicts, each explaining under four percent of variance.

A further consideration reinforces why single-gene copy-number tracking is more reliable here than bulk expression alone. Bulk RNA-seq measures a mixture of tumor, immune, and stromal cells, and *PTEN*-deleted tumors differ from *PTEN*-intact tumors in immune and stromal composition; part of any bulk transcriptomic difference across dosage tiers could therefore reflect shifts in microenvironmental composition rather than tumor-cell-intrinsic regulation. This confound does not affect the central analysis, because the copy number of *GLUD1* and *GOT1* is measured in the same tumor DNA that defines *PTEN* status and tracks it directly, whereas microenvironmental variation cannot create a gene-dosage correlation between physically linked loci. Where composition could plausibly have mattered, in the prostate-specific association, consensus purity estimates were included as a covariate and the association persisted. The concordance between the gene-level copy-number evidence and the expression pattern therefore argues against a microenvironmental explanation and for structural co-deletion. Beyond *PTEN*, the result is a specific instance of a general and under-appreciated problem in signature-based cancer genomics. Multi-gene expression or copy-number signatures are widely used to infer pathway activity from tumor profiles, yet when a signature contains genes that reside near a recurrently deleted driver, deletion of the driver mechanically alters the signature through loss of the linked passengers rather than through the biology the signature is meant to report. Canonical pathway gene sets used for enrichment scoring place *GLUD1* and *GOT1* in the same set as *GLS* and *GPT2* (KEGG alanine, aspartate and glutamate metabolism, MSigDB M17758, a member of the KEGG legacy subcollection), so *PTEN*-centered metabolic signatures are particularly exposed to this confounding. Our interval analysis shows the exposure is positional rather than pathway-specific: across 10q23-q24, co-deletion with *PTEN* was strongly and monotonically related to distance, and genes carrying no glutaminolysis annotation were indistinguishable from *GLUD1* and *GOT1* in this respect. The safeguard is straightforward and was applied here: decompose a signature by chromosomal location relative to the driver, and test the constituent genes’ copy number against the driver’s copy number, before interpreting an association as regulatory. For studies specifically aiming to measure transcriptional regulation of glutaminolysis in the context of *PTEN* loss, we suggest a chromosome-aware signature that excludes the 10q23-q24 passengers *GLUD1* and *GOT1* and relies on the off-chromosome-10 members (*GLS*, *SLC1A5*, *GPT2*), optionally supplemented with additional non-chromosome-10 glutaminolytic genes such as *GLS2* or *SLC38A1*. In the present data this reduced signature showed no decline with *PTEN* loss in most tumor types, indicating that it isolates regulatory signal from the structural artifact. The confound can be screened for in four steps, the first three of which require only data already in hand. (1) Stratify the signature genes by chromosomal position relative to the deletion driver. (2) Correlate each gene’s own copy number against the driver’s copy number, flagging any gene whose copy number tracks the driver as a candidate passenger. (3) Where mutation data allow, compare tumors carrying an inactivating driver mutation without deletion against copy-neutral wild-type tumors, since a passenger gene will respond to deletion but not to mutation alone. (4) Use independent functional data, such as CRISPR dependency screens, as an orthogonal check on whether the inferred pathway activity corresponds to a real cellular dependency, bearing in mind that dependency is itself context-dependent. We propose this as a screening workflow rather than a validated instrument. The principal implication is methodological and cautionary. Copy-number-derived or expression-derived gene signatures that include genes physically near a driver locus can produce associations that mimic coordinated regulation but arise from passenger co-deletion. *GLUD1* and *GOT1* sit within the commonly deleted 10q23-q24 interval, so any *PTEN*-centered metabolic signature that contains them will appear to track *PTEN* loss for structural reasons. Signature design should account for the chromosomal location of constituent genes relative to the driver under study, and associations of this kind should be tested against the copy-number of the individual genes, as done here.

The confound identified here is distinct from the therapeutic opportunity that has motivated the collateral-lethality literature. Passenger co-deletion is well established: homozygous deletion of a tumor suppressor routinely removes neighboring sequence, and this has been developed into a therapeutic strategy. Deletion of ENO1 alongside 1p36 creates a selective dependence on ENO2 in glioblastoma [30], deletion of malic enzyme 2 alongside *SMAD4* creates a comparable vulnerability in pancreatic cancer [31], and the principle has been formalized as collateral lethality, the most widely exploited instance of which is the co-deletion of *MTAP* alongside *CDKN2A* at 9p21 [32]. The 10q23 region around *PTEN* is recognized as one of the more deletion-prone loci in cancer. Whether the same measurement confound arises at other deletion-prone loci is a question the present data cannot settle, since the procedure we set out below would have to be applied locus by locus. That literature asks which collaterally lost gene can be exploited. The present result asks what happens when collaterally deleted genes are members of a curated pathway signature: their loss changes the signature score mechanically, and the result is indistinguishable from coordinated regulation of that pathway.

For clinical translation, these data argue against using *PTEN* loss as a stand-alone biomarker to select patients for glutaminase inhibition. The transcriptional signature is reduced for structural reasons, *GLS* dependence does not track *PTEN* dosage in cell lines, and clinical results with glutaminase inhibition have so far been disappointing [2], although the trials conducted were not selected on *PTEN* status and did not include the tumor types analyzed here, so they neither confirm nor contradict the present finding directly. Genuine glutamine dependence, where it exists, arises through other alterations and requires direct functional or metabolic selection [33].

Limitations include reliance on copy-number-defined dosage, which does not capture point mutations, promoter methylation, or structural variants; some functionally *PTEN*-deficient tumors will be classified as intact, biasing toward the null. Epigenetic *PTEN* silencing can itself have metabolic consequences distinct from copy-number loss, as demethylating agents that restore *PTEN* expression have been shown to normalize glucose metabolism in gastric cancer cells [34], underscoring that copy-number dosage captures only one route to *PTEN* inactivation. The glioblastoma cohort had too few intact tumors for a reliable contrast. Thresholded GISTIC calls can assign homozygous status within a broad low-level loss, while low tumor purity works in the opposite direction by masking true homozygous deletion; both are sources of misclassification in the dosage tiers. Neither bias appears to dominate here. Homozygous deletion frequencies run from 2.9 to 10.8 percent across thirteen cohorts, with prostate carcinoma the outlier at 17.4 percent, which is in the range reported for focal *PTEN* events rather than for arm-level loss; and median tumor purity is lower in homozygously deleted tumors in only four of the fourteen cohorts, so purity-driven masking, where it operates, biases these frequencies downward rather than upward. *GLS* is a weak dependency in most cell lines irrespective of *PTEN* status, so the DepMap result bounds rather than proves the absence of a functional dependency. Bulk RNA-seq is also subject to tumor-purity and stromal-dilution differences that can vary systematically across *PTEN* dosage tiers; we therefore relied on gene-level GISTIC copy-number data, which are measured in tumor DNA and unaffected by such admixture, to establish the co-deletion mechanism, and this is why the copy-number evidence rather than expression alone is treated as decisive. Bulk expression carries a compositional caveat that the copy-number evidence does not. *GLS* and *SLC1A5* are expressed in immune cells, and *PTEN*-deleted tumors differ from intact tumors in immune and stromal content, so part of any measured change in the off-chromosome-10 sub-signature across dosage tiers may reflect cellular composition rather than tumor-cell transcription. This does not bear on the co-deletion finding, which rests on copy number measured in tumor DNA, but it is a reason to treat the expression-level interpretation of those three genes with caution in the lineages where they move at all. The signature is a transcriptional proxy and does not measure glutamine flux; however, the concordant DepMap dependency result and the direct copy-number evidence for co-deletion together support the central conclusion. The strength of the analysis is that the mechanism was tested rather than assumed, and a signaling interpretation was excluded in favor of a structural one supported by gene-level copy-number data.

## Author contributions

Conceptualization: Abdelmuhsen M. Abusneina. Formal analysis: Abdelmuhsen M. Abusneina, Soad M. Elwirfli. Methodology: Abdelmuhsen M. Abusneina. Writing - original draft: Abdelmuhsen M. Abusneina. Writing - review and editing: Soad M. Elwirfli. Both authors read and approved the final manuscript.

## Data availability

All data analyzed in this study are publicly available and require no special permissions. TCGA PanCancer Atlas copy-number, expression, mutation and clinical data were obtained from the cBioPortal datahub (https://www.cbioportal.org), with mutation data retrieved through the cBioPortal REST API, and consensus tumor purity from the TCGA PanCanAtlas ABSOLUTE tables. DepMap CRISPR gene-effect and copy-number data were obtained from DepMap Public 26Q1 (https://depmap.org/portal/download). The minimal data set required to replicate the findings reported here, comprising tumor-level derived variables for all analyzed cohorts together with a data dictionary and the list of studies assessed but excluded, is deposited at figshare (DOI: 10.6084/m9.figshare.33716476). The complete analysis code is archived at Zenodo (DOI: 10.5281/zenodo.21195209) and mirrored at https://github.com/Abusneina/pten-glutaminolysis-codeletion. No new data were generated by this study.

## Supporting information

**S1 Fig. Kaplan-Meier overall survival by *PTEN* copy-number dosage in four representative tumor types.** (A) Uterine endometrial carcinoma. (B) Glioblastoma. (C) Prostate adenocarcinoma. (D) Breast carcinoma. Tumors are grouped as *PTEN* intact, hemizygous deletion or homozygous deletion, with group sizes given in each panel.

S2 Fig. Glutaminolysis signature by *PTEN* dosage in the eight tumor types not shown in Fig 2. S1 Table. Copy number of each signature gene correlated with *PTEN* copy number.

S2 Table. Expression of each signature gene correlated with *PTEN* loss severity, with Cliff’s delta and bootstrap confidence intervals for each dosage comparison.

S3 Table. Partial correlation of each signature gene with *PTEN* loss severity, controlling for tumor aneuploidy, for a 10q arm-level proxy, and for each gene’s own copy number, each separately.

S4 Table. *GLUD1* expression in *PTEN*-mutant copy-neutral versus wild-type and hemizygously deleted tumors, per tumor type, including the truncating-only sensitivity analysis and stratification by microsatellite instability.

S5 Table. Studies assessed but not analyzed, with *PTEN* copy-number tier counts and the reason for exclusion.

S6 Table. Co-deletion with *PTEN* for thirty-seven 10q23-q24 genes carrying no glutaminolysis annotation, with distance from *PTEN*.

