## Supplementary figures and images for "Passenger co-deletion confounds glutaminolysis signatures anchored on *PTEN* loss: a cautionary case for location-aware signature design"

### S1_Fig

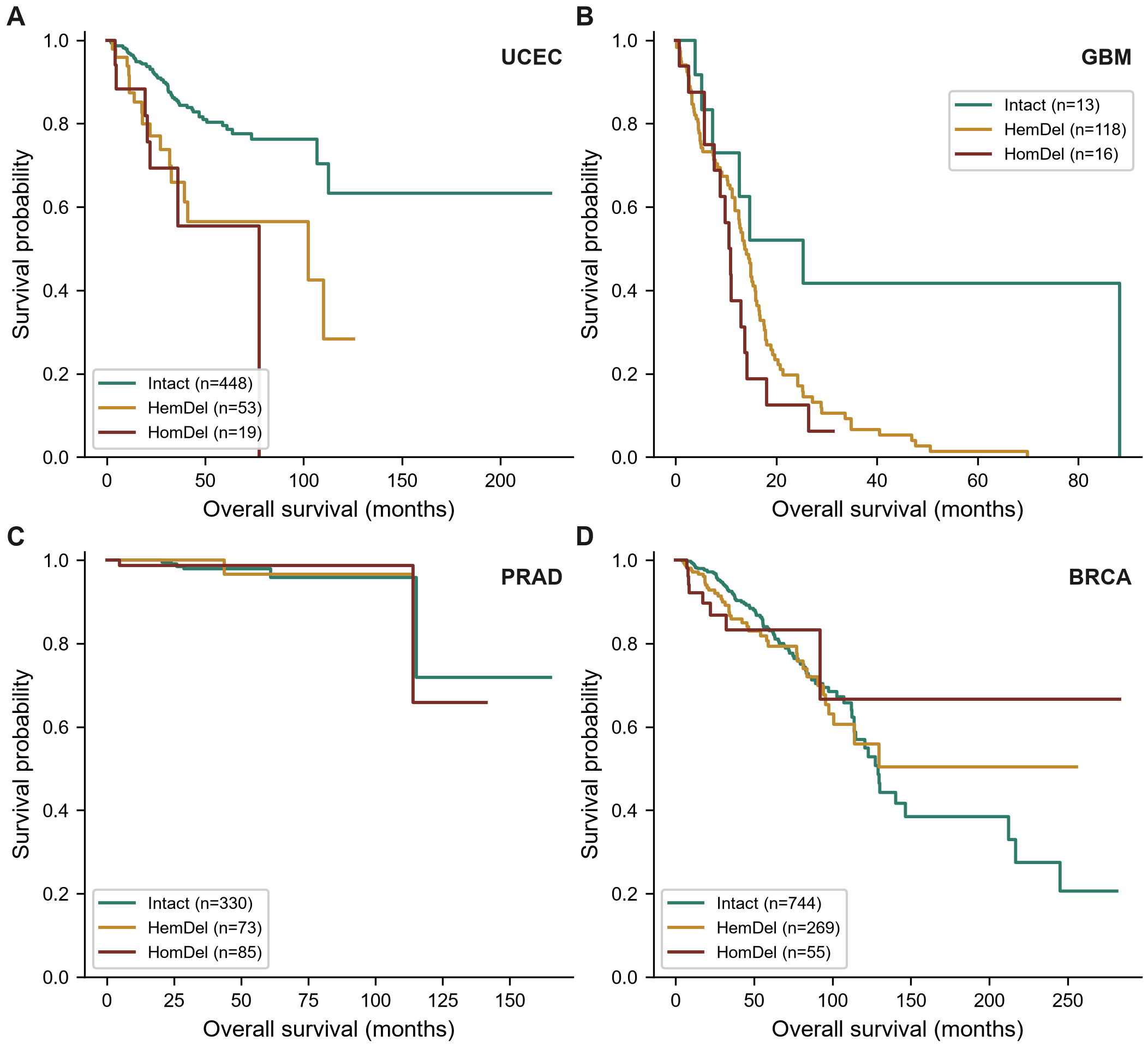

### S2_Fig

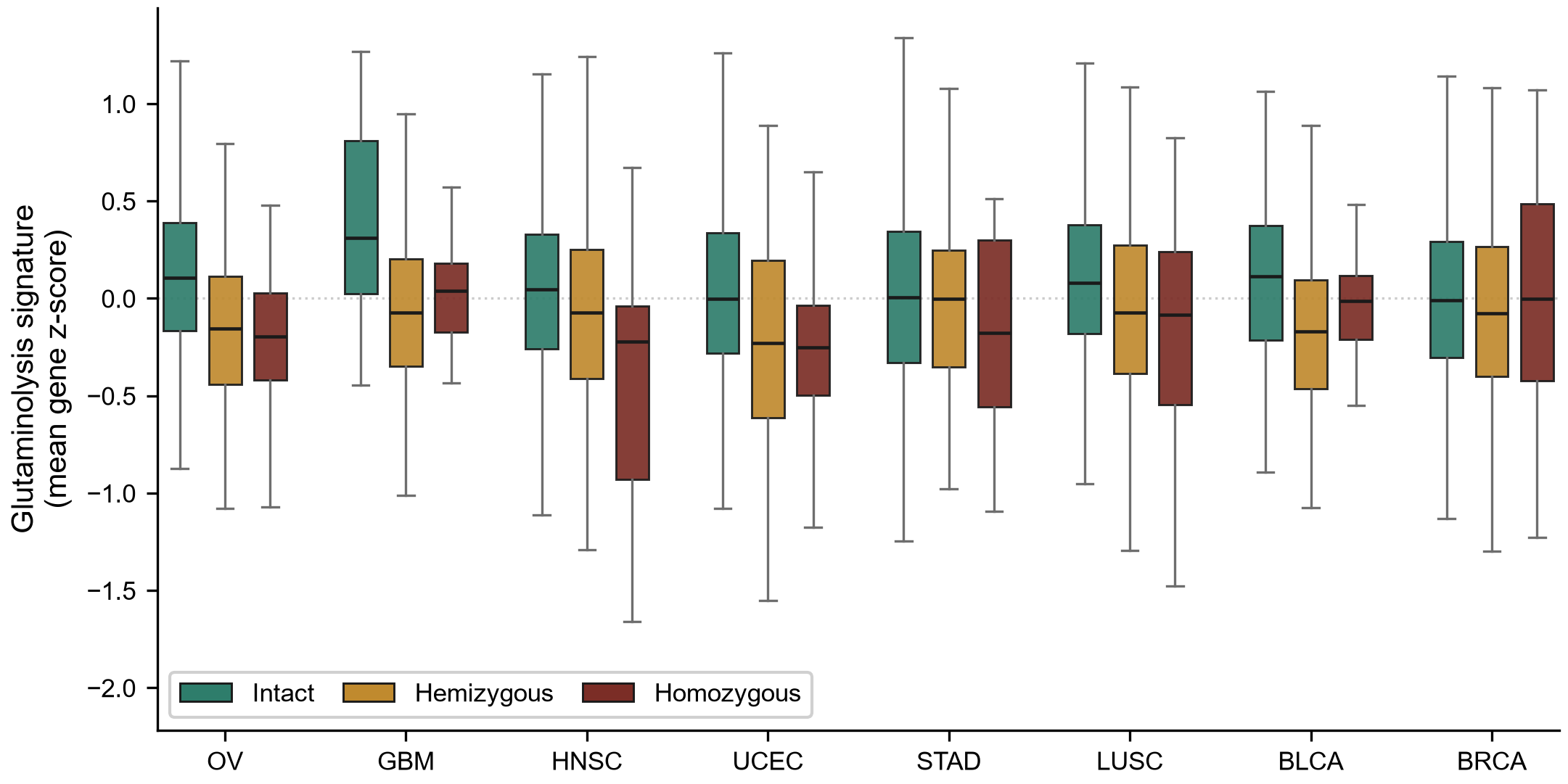
